# Skin Sensitisation Case Study: Comparison of Defined Approaches including OECD 497 Guidance

**DOI:** 10.1101/2024.02.12.579980

**Authors:** Pascal P. Ankli, Shaheena Parween, Béatrice Lopez, Pierre Daligaux, Tomaž Mohorič, Thomas Darde, Christophe Chesné, Nathan Stockman, Csaba Boglári, Amanda Y. Poon, Barry Hardy

## Abstract

Characterising known and new chemical compounds for skin sensitisation provides a basis for the development of safer products where ingredients are exposed to skin. By including new approaches, such as tiered testing strategies and integrated data analysis, it is possible to develop next generation products adhering to emerging regulations, scientific evidence and animal welfare principles. To ensure data integrity during such assessment the OECD provides characterisation guidelines and Defined Approaches (DAs) to uniform work-flows. In this study we developed and applied the integrated characterisation tool called «SaferSkin^™^» to compare the results of different DAs for eight compounds and included results obtained from current OECD guidance and emerging methods. We tested two compounds with unclear or indeterminate results with the SENS-IS assay to explore the value of the experiment in strengthening the weight of evidence and arriving at a clearer conclusion.

## Introduction

Skin sensitisation is defined as the molecular biological endpoint which describes the allergic response induced by repeated exposure of a chemical to the skin leading to Allergic Contact Dermatitis (ACD). (1) ACD thereby is characterised by a sensitisation event as a result of the activation of the immune response by an allergen and an elicitation event as a result of repeated exposure to the allergen. ACD causes an itchy rash on skin which can be very uncomfortable. ACD is induced by a wide range of ingredients from products which are exposed to skin including cosmetics, fragrances, natural products derived from plants and materials such as textiles and jewellery. (2) To ensure consumer safety and to develop safe and sustainable products it is essential to assess ingredients for their risk to cause skin sensitisation in concern of ACD.

Until now, assessment approaches mainly included methods such as human patch testing and the Lymphocyte Transformation Test (LTT). (3) Those methods have their drawbacks since clinical studies might be very time consuming and *in vitro* tests may not reflect the true response of a compound in living organisms. To evaluate compounds in concern of their safety efficiently, the Organisation for Economic Cooperation and Development (OECD) has elaborated Defined Approaches (DAs) and guidelines to assess chemicals and materials in form of tiered strategies. (4) Tiered strategies allow to evaluate chemicals and materials based on experimental needs in such a way that complex and time-consuming test methods are only then conducted when necessary. Otherwise, next generation machine learning and Artificial Intelligence (AI) computational approaches are used to evaluate and integrate already existing experimental data. This *in silico* strategy allows to characterise compounds by feature extraction and ontology computation in relation to molecular structures of known pharmacophores which show a pharmacological effect on molecular mechanisms which are involved in causing the phenotype. The *in silico* characterisation thereby is integrated with results which come from already existing *in vivo* data and existing data from clinical studies and thereby ensure the integrity of the approach. (5)

To evaluate chemicals and materials in concern of skin sensitisation, molecular Key Events (KEs) and Adverse Outcome Pathways (AOPs) have to be defined against which the chemicals and materials can be evaluated. The Adverse Outcome Pathways on skin sensitisation have been well described in the OECD 497 Guidance (4) and the OECD report on «The Adverse Outcome Pathway (AOP) for Skin sensitisation Initiated by Covalent Binding to Proteins». (6) Skin sensitisation thereby is characterised by a Molecular Initiating Event (MIE) e.g. a first Key Event (KE1) where chemicals covalently bind to partially negative skin proteins what leads to an activation of molecular mechanisms where several further KEs can be defined downstream in the pathway. The second KE (KE2) is defined such that inflammatory responses are induced in keratinocytes, which constitute the main cell type in epidermal skin tissue. (7) Altered gene expression, as a hallmark of this KE, further leads to impaired cell signalling. The third KE (KE3) includes the activation of dendritic cells and the presentation of chemokines and cytokines as biomarkers on their cell surface before T - cells get activated, what constitutes the fourth KE (KE4). The cascade of KEs ends with an adverse outcome which can be taken as the endpoint for skin sensitisation and which is characterised as Allergic Contact Dermatitis. (6) There exist specific assays on specific key events as visualised in Figure 1. Some of those assays have been individually validated by the OECD. The green highlighted assays are OECD TG 497 assays and the ones which are in **Black bold** were used for the SaferSkin evaluation.

**Fig. 1:**
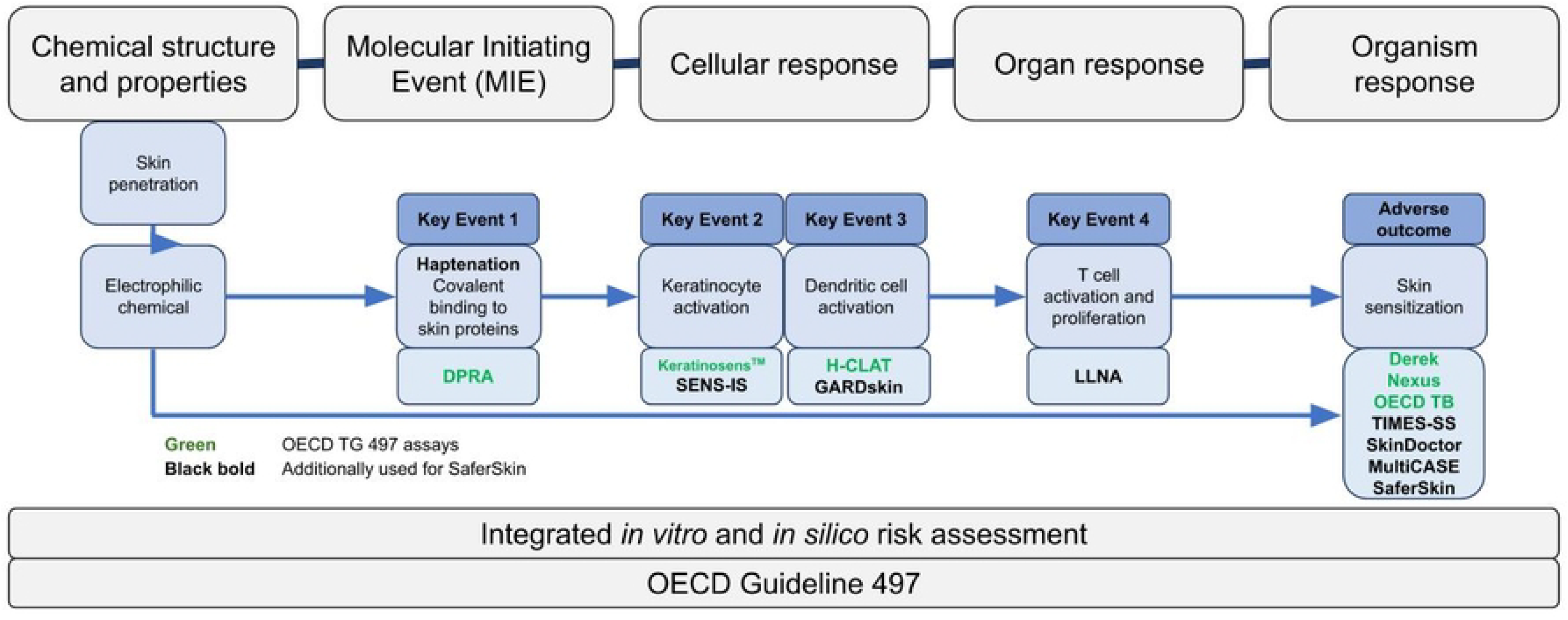
Available assays specific for different key events on skin sensitisation. Green highlighted assays = OECD TG 497 assays, **Black bold** = Assays additionally used in this study

To ensure the validity of results during the evaluation of such molecular events during the risk assessment process, standardised protocols and approaches have to be developed to ensure the integrity of experimental outcomes. Therefore, the OECD has elaborated such Defined Approaches (DAs) and Integrated Approaches to Testing and Assessment (IATA). (4),(8)

The OECD DAs thereby were structured such that experimental values have to be confirmed by a biological assay approach which is combined with an *in silico* modelling approach and consists of the following three methods:

1. **A «Two out of three» (2o3)** hazard identification approach based on KE1, KE2 and KE3.
2. Either an **integrated strategy (ITSv1)** based on *in chemico* (KE1) and *in vitro* (KE3) data as well as on Derek Nexus *in silico* predictions.
3. Or a modification of the **integrated strategy (ITSv2)** based on *in chemico* (KE1) and *in vitro* (KE3) data as well as on OECD QSAR Toolbox *in silico* predictions.

We chose ITSv2 over ITSv1 for the comparisons reported in this article.

During the evaluation and comparison of method results, it is preferred to include data from validated test methods having OECD test guidance, which in this case includes the Direct Peptide Reactivity Assay (DPRA) for KE1 (9), the Keratinosens^*™*^ Assay for KE2 (10) and the h-CLAT Assay for KE3. (4), (11)

Based on this background we developed a predictive approach following the OECD 497 guidelines and incorporated different tools into an integrated solution package called «SaferSkin». «SaferSkin» is one of many services from «SaferWorldbyDesign». «SaferWorldbyDesign» is a platform which was developed by Edelweiss Connect GmbH and builds a framework for risk assessment characterisations of all kinds of chemicals and materials. SaferWorldbyDesign interacts with customers and adapts its services for specific customer needs with a strong partner consortium as part of the solution packages. Figure 2 provides an overview over the work-flow during risk assessment within the solution packages at SaferWorldbyDesign.

**Fig. 2:**
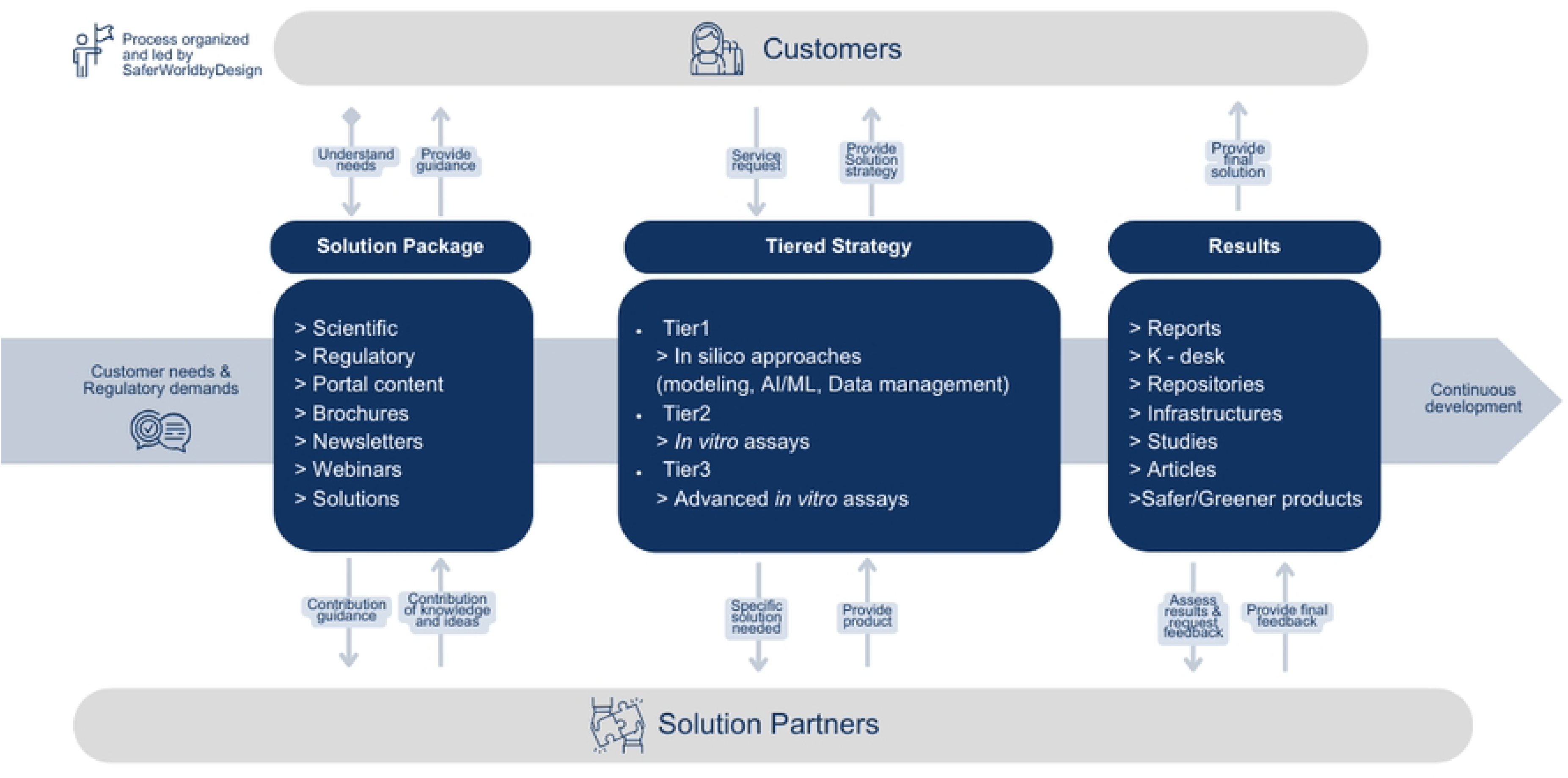
Work-flow of SaferWorldbyDesign risk assessment within a possible solution package, such as SaferSkin.

«SaferSkin», the solution package, includes the OECD required *in vitro* assays DPRA (9), Keratinosens (10) and h-CLAT (11), the OECD required *in silico* assays Derek Nexus (KREATiS) (8) and OECD Toolbox (12) in the 2o3 ITSv1 and ITSv2 approaches. Further assays in «SaferSkin» include the *in vitro* assays GARD (13) and SENS-IS (14) as well as the *in silico* tools including the SaferSkin application with its Bayesian network (15) and multiple regression model (16), the MultiCASE CASE Ultra Toolbox (17), SkinDoctor (18) and TIMES SS Globally Harmonized System (GHS) (19) which we offer in a multi-tier approach depending on customer needs. The results of the assessment process serve as a guidance for product developers to make their products safer and more sustainable to ensure consumer safety and to face consumer needs and expectations.

During assessment, results might be unclear. With the SENS-IS assay (14) we hereby present a method which we used as an additional approach where outcomes from the first and second tier approaches remained unclear.

During this study we evaluated the different elements of our tiered approach from «SaferSkin» in concern of its functionality during case studies on a variety of compounds. With this, we present a tool that allows us to evaluate compounds in concern of skin sensitisation in a fast, efficient and sustainable way.

## Methodologies

### Modelling and data analysis

The components of our assessment process consisted of:

- **The current OECD TG 497 guidance**, including 2o3 and ITSv2 DA as implemented within the OECD toolbox. (4)
- **The SaferSkin application**, our computational skin characterisation comparison tool, including Bayesian network (15) and multiple regression methods (16).
- **Assay result data** including DPRA (9), Keratinosens (10), h-CLAT (11), GARD (13) and **SENSIS (14)**.
- **The TIMES SS (GHS) model**, a skin metabolic simulator to underline assessment results. (19)
- **The SkinDoctor application**, which is a random forest model for binary classification of sensitisers versus non-sensitisers. (18)
- **The MultiCASE CASE Ultra Toolbox**, which is a QSAR software for modelling and predicting toxicity of chemicals. (17)
- **Reference data** based on already existing human patch (20) and Local Lymph Node Assay (LLNA) (21) data.

**2o3** includes the DPRA (KE1), Keratinosens (KE2) and h-CLAT (KE3) assays. In the DPRA assay a chemical is exposed to cysteines (Cys) and lysines (Lys) where the binding capacity of the test chemical to Cys and Lys is detected by HPLC analysis. In DPRA, a chemical is considered positive for skin sensitisation if the mean peptide depletion is ≥ 6.38 % or in the case of co-elution, cysteine-only depletion above 13.89 %. In the Keratinosens assay a gene expression profile is established for the AKR1C2 gene in keratinocytes, which is known to be over-expressed during skin sensitisation. Gene expression is measured by spectrophotometry using a luciferase target gene linkage approach. In the Keratinosens assay, compounds are considered positive when the luciferase activity (I_*max*_) ≥ 1.5 - fold the I_*max*_ compared to control below a defined concentration which does not significantly affect cell viability (i.e. c *<* 1000 *µ*M at which the cell viability is *>* 70 %). In the h-CLAT assay, antibodies against CD54 and CD86 are used to measure their expression on dendritic cells by flow cytometry. In the h-CLAT assay, chemicals are considered positive when their fluorescence intensity is ≥ 1.5-fold for CD86 or ≥ 2.0-fold for CD54 at any concentration with cell viability *>* 50 %. With this, compounds are either considered as positive for skin sensitisation or negative for skin sensitisation. (4)

If one of the assays results in a value close to the threshold, we characterise this compound to be a borderline value. Borderline compound values can still be considered if the other two assays out of the three (2o3) assays give a non-contradicting, clear result. Borderline Ranges (BR) for the assays are: (4)

- **DPRA BR:** Mean peptide depletion of 4.95 % to 8.32 %, Cys only depletion in the case of coelution with lysine: 10.56 % to 18.47 %.
- **Keratinosens BR:** I_*max*_: 1.35-fold to 1.67-fold.
- **h-CLAT BR:** Relative Fluorescence Intensity (RFI) CD54: 157 % - 255 %; RFI CD86: 122 % - 184%.

**ITSv2** takes the outcomes of the 2o3 approach into account and integrates it with the OECD QSAR *in silico* characterisation. Thereby, score values of either 0 or 1 are given for the *in silico* prediction and score values 0, 1, 2 or 3 for the DPRA and the h-CLAT assays. Those score values sum up to a total score of either 0, 1, 2, 3, 4, 5, 6 or 7 according to the United Nations Globally Harmonized System (UN GHS) to define the categories Not Classified (NC), In-Conclusive (IC), 1A (strong sensitisers) and 1B (other sensitisers). The criteria on what score is given to each compound are based on the assay outcomes and dependent on the Minimum Induction Threshold (MIT) for h-CLAT, the mean Cys and Lys depletion, the stand alone Cys depletion for the DPRA assay and the positive or negative outcome for the ITS approaches. (4) The scoring dependence is visualised in Table 1 for each of the approaches, respectively.

**Table 1:** OECD Scoring based on experimental outcome.

| Score | DPRA |  | h-CLAT | QSAR |
| --- | --- | --- | --- | --- |
| | Depletion [%]<br>Cys/Lys | Cys | MIT [$\mu\text{g/ml}^g$] | |
| **3** | $\geq 42,47$ | $\geq 98,24$ | $\leq 10$ | |
| **2** | $\geq 22,62,$ | $\geq 23,09,$ | $> 10,$ | |
| | $< 42,47$ | $< 98,24$ | $\leq 150$ | |
| **1** | $\geq 6,38,$ | $\geq 13,89,$ | $> 150,$ | |
| | $< 22,62$ | $< 23,09$ | $\leq 5000$ | + |
| **0** | $< 6,38$ | $< 13, 89$ | not calc. | – |

Based on the individual scores, total scores are evaluated including borderline outcomes. For this, a work-flow has been defined which clarifies under given outcomes how one should proceed and what results have to be taken into account for the final total score. If two assays are applicable for evaluation and the *in silico* prediction is in domain, then the final score is defined as the sum of the individual DPRA, h-CLAT and QSAR scores. If the *in silico* prediction is out of domain, the final score will be the sum of the individual DPRA and h-CLAT scores and if there is only one assay available the sum of the individual available assay and QSAR scores will give the final evaluation score. (4) The scoring procedure is summarised in Table 2.

**Table 2:** OECD standardization of ITS scoring.

| *Two assays applicable* |  | *One assay applicable* |
| --- | --- | --- |
| ***In silico* prediction in domain:** | ***In silico* prediction not in domain:** | ***In silico* prediction in domain:** |
| Sum of DPRA, h-CLAT and QSAR | Sum of DPRA and h-CLAT | Sum of applicable assay and QSAR |
| **Score: Prediction** | **Total score: ITS prediction** | **Total score: ITS prediction** |
| 6-7: UN GHS 1A | 6:UN GHS 1A | 3-4: UN GHS 1 (hazard) |
| 2-5: UN GHS 1B | 5:UN GHS 1 (hazard) | 2:UN GHS 1B |
| 0-1: NC | 2-4: UN GHS 1B | 0-1: Inconclusive |
|  | 1 :Inconclusive |  |
|  | 0 :NC |  |

**The OECD Toolbox** is an application which was developed by the OECD with the aim of providing an integrated *in silico* approach to assess the risk of chemicals being hazardous to all fields of industry. The toolbox works by identifying structural features of chemicals which then are correlated with specific modes of molecular action. Based on this ontology comparison, predictions are made for new compounds and evaluated against existing experimental data to underline the prediction outcome. Thereby, the OECD QSAR Toolbox offers three approaches: (12) A read-across approach where endpoint information is predicted by existing data of that endpoint coming from other compounds, a trend analysis approach where existing data is collected during a certain timespan that might be important for assays such as FISH, LC_50_ etc. and a Quantitative Structure-Activity Relationship (QSAR) approach which uses machine learning algorithms that learn from existing training data sets and where new data sets are evaluated based on the trained model. For our evaluations here we used the QSAR approach.

**The SaferSkin application** was developed by Edelweiss Connect to provide an easy-to-use prediction tool to evaluate chemicals as potential skin sensitisers with multiple model comparisons and reporting. It consists of a user input panel where the compound structure can be entered either as a Simplified Molecular Input Line Entry System (SMILE) structure or by drawing the molecular structure of interest in a drawing panel. Experimental values, such as protein binding capacity and solubility, are calculated by QSAR descriptors or may be directly entered as overriding experimental values. (12) Additionally, experimental *in vitro* assay results from DPRA, Keratinosens and h-CLAT assays can be entered for the Bayesian network as well as for the multiple regression models to be able to make a prediction and to proceed with the 2o3 scoring. (22) The Bayesian network is a graphical based computational approach to represent knowledge. The model links variables with probabilities to affect other variables which are interlinked with themselves. This allows to make predictions of new variables by the existing knowledge. (23) The Multiple Regression approach is based on independent variables with which the value of a dependent variable can be predicted. (24) With this data, a prediction can be made which gives you a comparison of the results for the Bayesian network calculation, the multiple regression approach as well as for the 2o3 evaluation, respectively, including a prediction of confidentiality indication. The results can be archived and shared in the form of a detailed report.

With those two models we have a strong predictive tool to predict skin sensitisation for known and new compounds. Nevertheless, by including more models, the predictions can be improved. Therefore, we also included the following models for our predictions:

- **The Genomic Allergen Rapid Detection (GARD) assay** which is based on the Nano string nCounter technology, which enables the assessment of up to 800 genes or 228 gene fusions in twelve samples in a single assay. With a Support Vector Machine (SVM) model, chemicals are classified into category 1A (strong sensitisers) or 1B (other sensitisers). The assay measures dendritic cell activation and hence we used it for KE3 when other outcomes were not applicable or inconclusive. (13)
- **The TIMES SS (GHS) model** which was developed by the Laboratory of mathematical chemistry in Bulgaria and which is a Skin Metabolic (SM) simulator based on empirical and theoretical knowledge. It considers non-enzymatic transformations, enzyme-mediated reactions and Protein Binding Reactions (PBR). (25)
- **The SkinDoctor application** is an *in silico* tool which is a random forest model for binary classification of sensitisers versus non-sensitisers. (18)
- **MultiCASE CASE Ultra Toolbox** is a QSAR software for modelling and predicting toxicity of chemicals for different types of applications. In our study we used the toolbox that was developed for predicting skin sensitisation. The tool provides predictions for ten models in total, including eye Draize, eye irritation, LLNA CAT2, LLNA CAT3, LLNA CAT4, Skin carcinogenicity, Skin corrosion, Skin irritation, Skin SENS LLNA and Skin SENS non LLNA. (17)
- **Reference Data** was used from databases containing already existing human patch assay data (20) and data from Local Lymph Node Assays (LLNA). (21)

**The SENS-IS assay** is a patented reconstructed human Episkin^®^ model assay which allows the testing of chemicals based on the analysis of the expression from a large consortium of different genes. The assay is robust, easily transferable, has high predictability and is reproducible. (14) The assay measures keratinocyte activation and hence was used for KE2 to underline the Keratinosens results. The assay is one of the new approach methods for evaluating skin sensitisation potential of ingredients and formulations. It is based on the measurement of gene expression of 61 skin irritation and sensitisation-related genes in the human 3D reconstructed epidermis model (Episkin^®^) upon exposure to the tested sample. Expression of these genes is quantified by a widely used RT/q-PCR technique. Based on the number of significantly induced genes, the tested sample can be classified as skin irritant or skin sensitiser. Thereby, the category 1B is further divided into weak (50 % compound concentration) and moderate (10 % compound concentration) outcomes and 1A is further divided into strong (1 % compound concentration) and extreme (0.1 % compound concentration) outcomes. (14) With this, we have a strong and valuable predictive tool which we can use as an additional approach for skin sensitisation risk assessment.

**In the case studies** we tested our *in silico* assessment approach with eight compounds: Farnesal, which is a flavouring agent and used as a food additive (26); Safranal, which gives Saffron its aroma. Safranal has a variety of pharmacological functions. It is an agonist at GABA_*A*_ receptors, it has high antioxidant activity, it shows cytotoxicity against cancer cells *in vitro* and acts as antidepressant (27); 2-Butoxyethyl acetate which is widely used for varnish and solvent for colour pigments (28); 3-(Diethylamino) propylamine, which is an additive in coatings and plastic manufacturing (29); Furil, which is used as starting material in chemical industry (30); Benzyl alcohol, which is mostly used as a solvent, as bacteriostatic and in the treatment of head lice (31); Squaric acid, which is used in the treatment of warts and alopecia (32) and ethyl (2E,4Z)-deca-2,4-dienoate, which is a plant metabolite and used as flavouring agent and as a fragrance. (33) To evaluate the compounds, we followed a standardised *in silico* procedure: First we started with the evaluation of data from DPRA assay to evaluate the mean peptide depletion for cysteine and lysine given in [%], where a compound is considered positive when it causes a peptide depletion ≥ 6.38 % or in the case of co-elution with lysine, cysteine-only depletion above 13.89 %. For the Keratinosens assay we used the data on the concentrations in [*µ*M] which lead to a 1.5-fold (KEC_1.5_), or 3-fold (KEC_3_) luciferase induction as well as the IC_50_ value which describes the concentration which effects a reduction of cellular viability by 50 %. With data on the h-CLAT assay we derived the concentrations for which a test compound induced a Relative Fluorescence Intensity (RFI) of either 150 e.g. 1.5-fold (EC_150_) or 200 e.g. 2.0-fold (EC_200_). The CV_75_ value describes the concentration showing 75 % of cell survival (25 % cytotoxicity). Based on the outcomes of those three assays, we classified the compounds for the 2o3 approach into either a skin sensitiser or skin nonsensitiser. We also scored the compounds based on the ITSv2 scoring approach according to Table 1 and assessed a total sensitisation score by implementing the result from the OECD toolbox characterisation, which has either score 1 for sensitiser results or 0 for non-sensitiser results. We further evaluated the compounds with the Bayesian network, the multiple regression, TIMES SS computations, the SkinDoctor application as well as the MultiCASE CASE Ultra Toolbox and finally compared the results with the outcomes when available from the SENS-IS assay, the GARD assay and the reference human patch and LLNA data to give a final prediction comparison.

In some cases, different approaches may reveal different results. When we faced contradictory results in our assessment process, as it was the case for squaric acid, or when the compound could not be assessed due to solubility problems as it was the case for ethyl (2E,4Z)-deca-2,4-dienoate, we used experimental SENS-IS assays as an additional tool to predict the sensitisation.

### The SENS-IS experimental assay

The SENS-IS assay takes into account 61 genes where 23 of those genes are biomarkers for skin irritation and 21 «SENS-IS»-genes as well as 17 «ARE»-genes are biomarkers for skin sensitisation. Skin irritation is caused by the chemical itself whereas sensitisation involves the activation of immune responses. (34) With this, the assay offers prediction not only for skin sensitisation but also for skin irritation. The assay approach starts with a solubility test where compounds are evaluated according to their solubility in PBS, Olive Oil (OO) and DMSO at 10 % and 50 % concentrations at RT and 37 ^*o*^C. 30 *µ*l of the substance is then exposed to a 3D human reconstructed epidermis (EpiSkin^®^) model for 15 min. and prepared for RT/q-PCR. The RT/q-PCR measurement is done with SYBRGREEN^®^, specific primers and Glucoronidase *β, β*2 microglobulin and Nono «non-POU domain containing octamer-binding» as housekeeping genes using the LightCycler 480 software. The mRNA content for each gene of interest is then normalised to the mean mRNA content of the 3 house-keeping genes and calculated as the expression level for the test compound divided by the expression level for the vehicle controls (DE) as visualized in Equation 1.

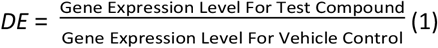

#### Test item classification

A test item is classified as irritant if at least 16 out of 23 genes of the «IRRITATION» group are significantly over-expressed.

A test item is classified as a sensitiser if at least seven out of 17 genes of the «ARE» group, and/or seven out of 21 genes in the «SENS-IS» group are significantly over-expressed.

The Cp value of the HSP90AA1 gene thereby must be ≤ 21. If more than 20 genes are over-expressed in the «IRRITATION» set of genes for a given concentration, the result is classified as false positive to take into account non-specific genes up-regulation that could be due to cell stress.

Moreover, the results obtained with the different concentrations allow the classification of the test item according to the lowest concentration that gives a positive result (DE *>* 1.25). Thus, a test item is classified in:

- Category 1A: Strong to extreme skin sensitiser, when a positive result is obtained at concentrations of 1 % and at 0.1 %.
- Category 1B: Weak to moderate sensitiser, when a positive result is obtained at concentrations of 10 % and/or 50 %.

A test item is classified as a non-sensitiser when negative results are observed at 100 % and at all other analysed concentrations. At least two independent experiments (repetitions) were performed in order to obtain two concordant conclusions. If three repetitions should be performed for a given concentration, a majority of positive results (2o3) must be obtained so that the final outcome is positive, otherwise the final outcome is negative.

## Results

### *In silico* results

For the eight compounds which we have tested, the results from the assay studies are listed in Table 3. The ranking for the 2o3 and ITSv2 approach, which includes the OECD Toolbox results, are summarised and compared in Table 4. A summary of all results is presented in Table 5.

**Table 3:**
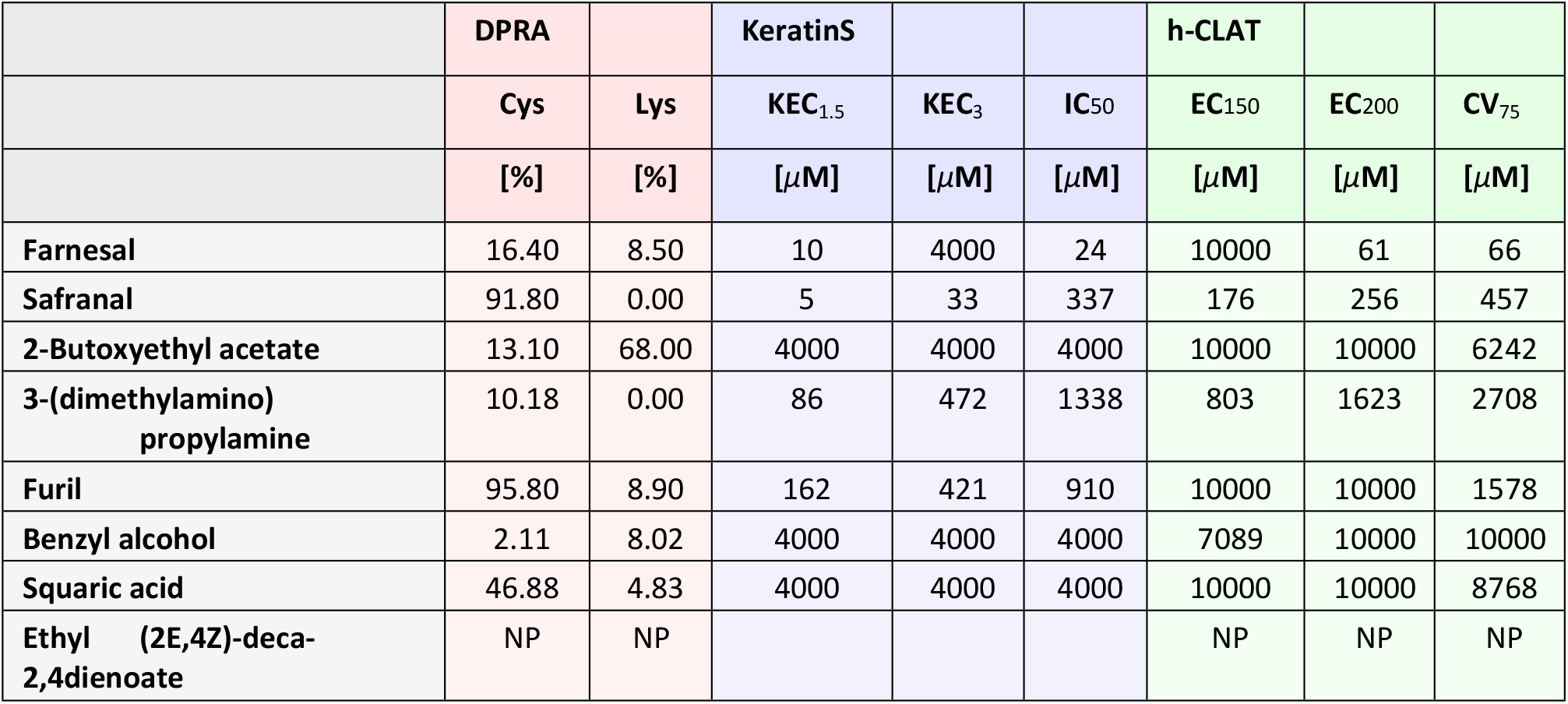
Summary of results from DPRA, KeratinoSens^™^ (KeratinS) and h-CLAT assays. For DPRA, the mean peptide depletion for Cys and Lys are given in [%]. For the KeratinoSens^™^ assay, the concentrations in [µM] are given which result in a 1.5 fold (KEC_1.5_) or 3-fold (KEC_3_) luciferase induction. The IC_50_ value describes the concentration which effects a reduction of cellular viability by 50 %. [10] For the h-CLAT assay, the concentrations are given at which the test chemicals induced a Relative Fluorescence Intensity (RFI) of either 150 e.g. 1.5-fold (EC_150_) or 200 e.g. 2-fold (EC_200_). The CV_75_ value describes the concentration showing 75 % of cell survival (25 % cytotoxicity. NP = Not Possible, empty fields = experiment not conducted.

|  | DPRA |  | KeratinS |  |  | h-CLAT |  |  |
| --- | --- | --- | --- | --- | --- | --- | --- | --- |
|  | Cys | Lys | KEC<sub>1.5</sub> | KEC<sub>3</sub> | IC<sub>50</sub> | EC<sub>150</sub> | EC<sub>200</sub> | CV<sub>75</sub> |
| | [%] | [%] | [μM] | [μM] | [μM] | [μM] | [μM] | [μM] |
| Farnesal | 16.40 | 8.50 | 10 | 4000 | 24 | 10000 | 61 | 66 |
| Safranal | 91.80 | 0.00 | 5 | 33 | 337 | 176 | 256 | 457 |
| 2-Butoxyethyl acetate | 13.10 | 68.00 | 4000 | 4000 | 4000 | 10000 | 10000 | 6242 |
| 3-(dimethylamino) propylamine | 10.18 | 0.00 | 86 | 472 | 1338 | 803 | 1623 | 2708 |
| Furil | 95.80 | 8.90 | 162 | 421 | 910 | 10000 | 10000 | 1578 |
| Benzyl alcohol | 2.11 | 8.02 | 4000 | 4000 | 4000 | 7089 | 10000 | 10000 |
| Squaric acid | 46.88 | 4.83 | 4000 | 4000 | 4000 | 10000 | 10000 | 8768 |
| Ethyl (2E,4Z)-deca-2,4-dienoate | NP | NP |  |  |  | NP | NP | NP |

**Table 4:** Ranking of results from DPRA, KeratinoSens^™^ (KeratinS), h-CLAT assay and the OECD Toolbox for the ITSv2 approach. According to Table 1 and Table 2. BL = BorderLine results; IC = InConclusive; + = positive for skin sensitisation; - = negative for skin sensitisation; empty fields = experiment not conducted.

|  | DPRA | KeratinS | h-CLAT | 2o3 | DPRA | h-CLAT | O TB | ITSv2 |
| --- | --- | --- | --- | --- | --- | --- | --- | --- |
|  |  |  |  | Score |  |  |  | Score |
| Farnesal | + | + | + | + | 1 | 2 | 1 | 4 |
| Safranal | + | + | + | + | 3 | 2 | 1 | 6 |
| 2-Butoxyethyl acetate | + | - | - | - | 0 | 0 | 0 | 0 |
| 3-(dimethylamino) propylamine | BL | + | + | + | 0 | 2 | 1 | 3 |
| Furil | + | + | BL | + | 3 | 1 | 1 | 5 |
| Benzyl alcohol | - | - | + | - | 0 | 1 | 1 | 2 |
| Squaric acid | + | - | - | - | 2 | 0 | 1 | 3 |
| Ethyl (2E,4Z)-deca-2,4-dienoate | IC |  | IC | IC | IC | IC | 1 | IC |

**Table 5:** Summary and comparison of the results from our Skin sensitisation assessment approach. **SSkin =** SaferSkin application, **OECD =** OECD Toolbox, **Ref. =** Reference Data, **SENS=** SENS-IS assay, **GAR =** GARD assay, **TIM =** TIMES SS model, **SD =** SkinDoctor application, **MC =** MultiCASE CASE Ultra toolbox, **Bay =** Bayesian network, **MR =** Multiple regression, **2o3 =** 2o3 approach, **ITS =** ITSv2, **LLNA =** LLNA reference data, **Hum =** Human patch reference data, NA = Not Available, NC = Not Classified, IC = InConclusive, NP = Not Possible, 1A = Strong skin sensitiser, 1B = Other skin sensitiser, W = Weak sensitiser, M = Moderate sensitisers, S = Strong sensitisers.

|  | SSkin |  | OECD |  | Ref. |  | SENS | GAR | TIM | SD | MC |
| --- | --- | --- | --- | --- | --- | --- | --- | --- | --- | --- | --- |
|  | Bay | MR | 2o3 | ITS | LLNA | Hum |  |  |  |  |  |
| Farnesal | M | M | + | 1B | 1B | NA | W | NA | + | + | + |
| Safranal | M | M | + | 1A | 1B | 1A | NA | NA | + | + | + |
| 2-Butoxyethylac. | - | - | - | NC | NC | NA | NA | NA | - | - | + |
| 3-(dimethylamino) propylamine | M | - | + | 1B | 1B | 1A | M | + | + | + | + |
| Furil | - | M | + | 1B | NC | NA | NA | NA | - | - | - |
| Benzyl alcohol | - | - | - | 1B | NC | 1B | W | + | - | - | + |
| Squaric acid | M | - | - | 1B | NA | NA | - | NA | + | + | + |
| Ethyl (2E,4Z)-deca-2,4-dienoate | NP | NP | NP | IC | NA | NA | S | NA | NA | - | + |

For farnesal the result is distinct. All three assays, the DPRA, Keratinosens and h-CLAT assay give positive results for sensitisation. Hence, the 2o3 prediction evaluates the compound as sensitiser. DPRA shows 8.5 % protein depletion for lysine and 16.40 % protein depletion for cysteine which leads to a score value of 1 (Table 1). The h-CLAT assay values result in a score 2 ranking. Also, the OECD Toolbox characterises farnesal as sensitiser what leads to a score 1. The summation of the individual scores results in a total ITSv2 score of 4 for farnesal. At least two assays were applicable and the *in silico* approach was in domain so that the final ranking is the result of the sum of the DPRA, h-CLAT and QSAR ranking what leads to a final score value of 4 (GHS 1B) what is described as a weak or moderate sensitiser. This result is confirmed by the Bayesian network, the multiple regression, SENS-IS, TIMES, the SkinDoctor application as well as the MultiCASE CASE Ultra Toolbox.

The same holds for safranal. All approaches characterise the compound as sensitiser. The Bayesian network and the multiple regression characterise safranal as a moderate sensitiser while the ITSv2 identifies it as a strong sensitiser according to human patch reference data.

2-butoxyethyl acetate gives an almost clear result. All approaches, except the MultiCASE CASE Ultra approach, identify the compound as a non-sensitiser. The MultiCASE CASE Ultra Toolbox characterises the compound as a sensitiser.

For 3-(dimethylamino) propylamine the characterisation values the compound either as a moderate or as a strong sensitiser. Only the multiple regression approach characterises the compound as a nonsensitiser, which is a troubling conflicting answer for this algorithm in this case.

For furil we have a more unclear situation as for 3-(dimethylamino) propylamine. The multiple regression agrees with 2o3 and ITSv2 to characterise the compound as a sensitiser. Now it is the Bayesian network which values this compound as a non-sensitiser according to the results which we get with the TIMES, the SkinDoctor and the MultiCASE CASE Ultra computational approaches.

For benzyl alcohol the result looks even more surprising. The ITSv2 characterises the compound as a sensitiser (but with a low score of 2), while the 2o3, Bayesian network and multiple regression all value the compound as a non-sensitiser. Also, the SkinDoctor application gives a negative result. When evaluating the compound with the SENS-IS and GARD assay, MultiCASE results and human patch data, the positive ITSv2 result is confirmed characterising benzyl alcohol as a weak/moderate sensitiser.

For squaric acid the results are also unclear. While the ITSv2 and the Bayesian network, TIMES, the SkinDoctor application and the MultiCASE CASE Ultra Toolbox characterise the compound as a sensitiser, the multiple regression and the 2o3 approach values it as a non-sensitiser. We hence selected this compound for SENS-IS experimental work for this case study.

For ethyl (2E,4Z)-deca-2,4-dienoate we cannot evaluate sensitisation. Phase separation occurred during the DPRA assay and for the h-CLAT assay the water solubility was too low to make a prediction (log(K_*ow*_) *>* 3.5). Since 2 out of the 3 assays were not conclusive the 2o3 approach was scored as inconclusive and leading to an ITSv2 score of 1, the ITSv2 score was hence inconclusive. While the SkinDoctor values the compound as a non-sensitiser, the MultiCASE toolbox characterises it as a sensitiser. The SkinDoctor application evaluates the compound as a non-sensitiser and the MultiCASE CASE Ultra characterises it as a skin sensitiser. Hence, we face a contradicting result with those predictions. We selected this second compound for SENS-IS experimental work for this study.

#### Metabolism

For the two compounds squaric acid and ethyl (2E,4Z)-deca-2,4-dienoate, where results were unclear, we included a metabolism analysis which we did with the OECD Toolbox, taking 3-(dimethylamino)propylamine as a third reference compound for this analysis. This analysis tool predicts possible metabolites derived from the test compound and summarises risk assessment results.

We got the following results:

#### 3-(dimethylamino)propylamine

- No alert found; 11 metabolites predicted by skin metabolism simulator and no metabolite in Autoxidation simulator in QSAR TB.
- For 5 out of 11 metabolites - protein binding alert found (Schiff base formation).
- Skin sensitisation data present for two metabolites (one sensitiser, one non-sensitiser).
- Using the skin permeability profiler (QSAR TB), high permeability is predicted for the parent compound and all metabolites. LogKp: 7.15 ^*<u>cm</u>*^_*s*_ (SwissADME).
- No CYP Inhibition (CYP1A2, CYP2C9, CYP2D6, CYP3A4).

#### Squaric acid

- Two metabolites predicted by skin metabolism simulator.
- Protein binding alert found (Schiff base formation).
- Two metabolites predicted by skin metabolism simulator in QSAR TB and one by autoxidation simulator (metabolite 1 is common).
- Protein binding alert found for both metabolites (Schiff base formation).
- Skin sensitisation data not present for both metabolites.
- High skin permeability predicted using toolbox profiler. LogKp: 6.58 ^*<u>cm</u>*^_*s*_ (SwissADME).
- CYP inhibition (CYP1A2, SwissADME).

#### Ethyl (2E,4Z)-deca-2,4-dienoate

- For parent compound - Protein binding alert found (Michaels addition).
- 13 metabolites predicted by autoxidation and skin metabolism simulator in QSAR TB.
- Protein binding alert found for ten out of the 13 metabolites (Michael addition - eight, Schiff base formation - two).
- Skin sensitisation data present for three metabolites - Non sensitiser in most assays, weak sensitiser in GPMT (One metabolite).
- High skin permeability predicted for the parent compound and nine out of the 13 metabolites and moderate skin permeability for four out of the 13 metabolites. LogKp: 4.65 ^*<u>cm</u>*^_*s*_ (swissADME).
- No CYP inhibition.

### The SENS-IS experimental results

Squaric acid was soluble in DMSO at 10 and 50 %

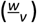 after heating the solution to 37 ^*o*^C.

Ethyl (2E,4Z)-deca-2,4-dienoate was soluble in olive oil and in DMSO at 10 % and 50 % 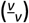 at Room Temperature (RT).

For **squaric acid** we get the results as summarised in Table 6.

**Table 6:** Results for squaric acid. For experiments where pure test compound (100 %) was applied to the epidermis model, 30 mg of solid test compound were put on the epidermis model and recovered by adding 30 µl of DMSO. Irrit. = Irritation, Sensit. = sensitisation.

| Squaric acid | Experiment 1 |  |  |  | Experiment 2 |  | Experiment 3 |
| --- | --- | --- | --- | --- | --- | --- | --- |
| Gene group | 100% | DMSO 50% | DMSO 10% | DMSO 1% | 100% | DMSO 50% | 100% |
| Irrit. | 17 | 12 | 21 | 12 | 16 | 6 | 20 |
| SENS-IS | 8 | 4 | 5 | 5 | 3 | 0 | 5 |
| ARE | 3 | 2 | 2 | 3 | 1 | 2 | 3 |
| Cp[HSP 90AA1] | 19.0 | 18.3 | 19.3 | 18.3 | 18.0 | 18.0 | 18.7 |
| Conclusion |  |  |  |  |  |  |  |
| Irrit. | Positive | Negative | Negative | Negative | Positive | Negative | Positive |
| Sensit. | Positive | Negative | Negative | Negative | Negative | Negative | Negative |

In the first experiment, squaric acid induced 8 genes in the «SENS-IS» gene group when it was tested at 100 % (pure test compound spread on the epidermis) and less than 7 genes in the «SENS-IS» and «ARE» gene groups when it was tested at 1, 10 and 50 % 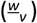 in DMSO.

In the second experiment, squaric acid induced less than 7 genes in the «SENS-IS» and «ARE» gene groups when it was tested at 50 % 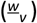 in DMSO and 100 % (pure test compound spread on the epidermis).

In the third experiment, squaric acid induced less than 7 genes in the «SENS-IS» and «ARE» gene groups when it was tested at 100 % (pure test compound spread on the epidermis).

Squaric acid gave negative results when it was tested in dilution at 1 % and 10 % in DMSO (one experiment), at 50 % in DMSO (two experiments) and 100 % in two experiments out of three and hence is validated as non-sensitiser.

**For ethyl (2E**,**4Z)-deca-2**,**4-dienoate** we get the results as summarised in Table 7.

**Table 7:** Results for Ethyl (2E,4Z)-deca-2,4-dienoate. OO = Olive Oil. Irrit. = Irritation, Sensit. = sensitisation.

| Decadienoate | Experiment 1 |  |  | Experiment 2 |  | Experiment 3 |  |
| --- | --- | --- | --- | --- | --- | --- | --- |
| Gene group | DMSO 10% | DMSO 1% | OO 10% | DMSO 10% | DMSO 1% | DMSO 1% | DMSO 0.1% |
| Irrit. | 18 | 5 | 0 | 16 | 11 | 10 | 7 |
| SENS-IS | 4 | 1 | 1 | 6 | 3 | 10 | 6 |
| ARE | 9 | 7 | 0 | 8 | 5 | 9 | 3 |
| Cp[HSP 90AA1] | 17.5 | 17.8 | 19.0 | 17.2 | 17.6 | 17.6 | 18.1 |
| Conclusion |  |  |  |  |  |  |  |
| Irrit. | Positive | Negative | Negative | Positive | Negative | Negative | Negative |
| Sensit. | Positive | Positive | Negative | Positive | Negative | Positive | Negative |

In the first experiment, ethyl (2E,4Z)-deca-2,4dienoate induced 7 genes or more in the «ARE» gene group when it was tested at 1 and 10 % 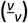 in DMSO and less than 7 genes in the »SENS-IS» and «ARE» gene groups when it was tested at 10 % 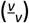 in olive oil.

In the second experiment, ethyl (2E,4Z)-deca-2,4dienoate induced eight genes in the «ARE» gene group when it was tested at 10 % 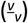 in DMSO and less than 7 genes in the »SENS-IS» and «ARE» gene groups when it was incubated at 1 % 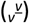 in DMSO.

In the third experiment ethyl (2E,4Z)-deca-2,4dienoate induced 9 and 10 genes in the «ARE» and «SENS-IS» gene group respectively when it was tested at 10 % 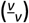 in DMSO. Less than 7 genes were overexpressed in both groups of sensitising genes when the test compound was tested at 0.1 % 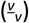 in DMSO. Ethyl (2E,4Z)-deca-2,4-dienoate gave positive results when tested at 1 % and 10 % in DMSO in two experiments out of three and hence is validated as a strong sensitiser of category 1A.

## Discussion

### *In silico* results

As we can draw from the results which we have obtained during the SaferSkin evaluation for our eight compounds, it is key to predict the biological behaviour of chemicals by a variety of different characterisation tools in an integrated approach to be able to receive meaningful results. Single evaluation tools might lead to misleading results which have to be verified and approved by further approaches. As we see from our studies, it is not only *in silico* computations which lead to contradicting results but also the *in vitro* assays for specific key events on biological endpoints may have uncertainties or borderline results.

To understand the cause of such contradicting results and to thereby improve the accuracy of predictions, Natsch et al. (35) proposes to take the following aspects into account:

1. Differences in prediction accuracy across potency classes for chemicals, where Natsch et al. suggest to more closely evaluate the reaction mechanisms of specific chemicals to correct under-predictions.
2. Different parameters which contribute to prediction in mechanistic domains, where they suggest developing more specific testing schemes for mechanistic domains in which prediction currently is insufficient. (36)
3. Different parameters which describe human and LLNA potency, which has to be considered during risk assessment to improve predictability.
4. Global versus domain models where a difference between domain prediction and actual *in vitro* results can be observed.
5. Insufficient statistical and mathematical approaches, where improving currently existing models and by using a variety of different available models may increase the prediction outcome. Also, by integrating bigger datasets, misclassification can be minimised.
6. Data redundancy, meaning that when three assays for three different key events of the same endpoint give the same result, taking all three outcomes for further data analysis might not increase the predictability although this might be expected by the risk assessor.

Taking those factors into account, contradicting results could be resolved by a weight of evidence approach to provide a conclusive prediction based on additional lines of evidence for given compounds by addressing the points above. (8),(37)

While Natsch et al. are suggesting improvements mainly for *in silico* methods, we must not forget that misprediction can also be caused by factors which come up much earlier in the prediction process. As Graudejus et al. (38) describe it in their paper, predictions that are based on *in vitro* results might not reflect how a compound is really affecting an organism *in vivo* for instance. While an *in vitro* result might be negative for skin sensitisation, a human patch test might reveal a positive result due to those differences. Not only between *in vivo* and *in vitro* approaches but even between single individuals of the same organismic kind there might exist variations due to genetic predisposition of an individual in concern of exposure to a compound. (39) Considering all those factors makes risk assessment a very complex task where all possibilities have to be taken into account to be able to make correct predictions.

As we can draw from our investigation there also might be solubility issues as we have faced it for the compound ethyl (2E,4Z)-deca-2,4-dienoate. Looking at the OECD guidelines OECD Guideline 497 (4), the OECD Guideline 442C for the DPRA assay (9), the OECD Guideline 442D for the Keratinosens assay (10) and the OECD Guideline 442E for the h-CLAT assay (11), solvents are commonly used such as listed below:

- **DPRA assay:** Acetonitrile, water, acetonitrile:water 1:1, isopropanol, acetone, acetone:acetonitrile 1:1. Also other solvents can be used as long as they do not have an impact on the stability of the peptide as monitored with reference controls. DMSO can only be used as a last resort and in minimal amounts. (9)
- **Keratinosens assay:** DMSO. If test chemicals are not soluble in DMSO, they can also be dissolved in water or culture medium. Also other solvents can be used with rational scientific arguments. (10)
- **h-CLAT assay:** Saline or medium as a first priority and DMSO as a second option. Other solvents can be used under reasonable scientific argumentation. (11)

When studying the article about the role of DMSO in cell mechanisms from Moskot et al. (40) we must assume that organic solvents not only change the chemical environment of a test compound and with it its reactivity but that the solvent also has direct effect on cell mechanisms which are not directly related to the test compound and hence might change an experimental outcome. Furthermore, if a test compound is not soluble in water and diluted in DMSO for instance, diluting it back into water based medium by a 25-fold, as it is suggested for the Keratinosens assay, it further rises the question of how effective the dilution of a test compound in DMSO is when changing the chemical environment back to water afterwards where the compound might fall out as a solid again and thereby change its effect onto the cell.

It is clear that for certain assays it is a challenge to dissolve test compounds in water-based solvents. Nevertheless, we raise concern about the procedure of dissolving compounds in non-water-based solvents and suggest that assays should be developed which mimic the *in vivo* behaviour of compounds as best as possible and to include solvent considerations in computational models to avoid mispredictions due to solvent-issues.

Adapting assays to *in vivo* conditions, developing new assays which mimic *in vivo* behaviour of chemicals better and including more experimental variables in computational models might hence increase the predictability of molecular endpoints better in the future.

With our study we evaluated different characterisation tools for our eight compounds and thereby show the importance of evaluating chemicals by a variety of different approaches to be able to receive an overall valuable result as we do for SaferSkin and suggest possible solutions to improve the characterisation work-flow for future approaches.

### SENS-IS experimental results

The strength of the SENS-IS assay lies in the genebased evaluation approach where already small changes in gene expression can be detected and where skin sensitisation is measured on the basis of molecular cell mechanisms.

The SENS-IS work reported here gave us results where previous methods had failed or where we faced contradicting results. Therefore, we show how important it is to integrate different characterisation tools to evaluate compounds for risk assessment and to not just rely on one approach.

## Conclusions

With this study we draw attention to the importance of integrated approaches during risk assessment of chemicals and materials by defined approaches and examine solutions to this task for skin sensitisation as provided by the SaferSkin application. We show that some methods might lead to unclear results and highlight that in such cases it is essential to underlie obtained results by further techniques such as the SENS-IS assay to clarify prediction outcomes.

## Supplements

### Links

- **Explore SaferWorldbyDesign including all mentioned assays:** [SaferWorldbyDesign](https://saferworldbydesign.com/).
- **Try out the SaferSkin Application:** [SaferSkin Application](https://saferworldbydesign.com/saferskin/in-silico/skin-sensitization-app/app/).
- **Try out the SkinDoctor application:** [SkinDoctor application](https://nerdd.univie.ac.at/skinDoctorII/).
- **Get the OECD Toolbox:** [OECD Toolbox](https://www.oecd.org/chemicalsafety/risk-assessment/oecd-qsar-toolbox.htm).
- **Get the MultiCASE CASE Ultra Toolbox:** [MultiCASE CASE Ultra Toolbox](https://multicase.com/case-ultra).
- **To get further information, please contact:** Edelweiss Connect GmbH.

